# Correlative humoral and cellular immunity to genetically attenuated malaria parasites in humans

**DOI:** 10.1101/2024.11.11.622940

**Authors:** Emil Colstrup, Rie Nakajima, Jelte M. M. Krol, Olivia A. C. Lamers, Rafael R. de Assis, Aarti Jain, Algis Jasinskas, Eva Iliopoulou, Helena M. de Bes-Roeleveld, Blandine M. D. Franke-Fayard, Meta Roestenberg, Philip L. Felgner, Rajagopal Murugan

## Abstract

Malaria caused by *Plasmodium falciparum* remains one of the major infectious diseases with a high burden in Sub-Saharan Africa. In spite of the advancements made in vaccine development and implementation in endemic countries, sterile and durable protection has not been achieved. Recently, we have shown the superior protective capacity of whole sporozoites attenuated to arrest late but not early during the liver stage development in a controlled human malaria infection study. Here we report the breadth of antigens targeted by hitherto understudied parasite liver stage immunity and convey the coherence between humoral and cellular immunity observed in our clinical study. Our findings uncover the underlying immunogenic differences between early- and late-liver stage arresting parasites and identify key liver-stage antigens for future vaccine development focused on inducing sterile immunity to malaria.

## Introduction

Malaria is one of the major life-threatening infectious diseases worldwide, causing 608,000 deaths and 249 million infections in 2023 (1). Among the five human infective species, *Plasmodium falciparum* (Pf) remains the cause of highest disease morbidity and mortality, particularly in low- and middle-income countries in Sub-Saharan Africa. Despite major efforts on disease control and vaccine development, reduction in malaria cases has stalled in recent years, challenging WHO global health and disease eradication programs. Currently, there are only two WHO-approved malaria vaccines available for clinical use: RTS,S/AS01 (Mosquirix) and R21/Matrix-M (2-4). While the vaccines significantly reduce the incidence of clinical malaria, the protective immunity tends to wane over time (5, 6).

Within 1-2 h post-infected mosquito bite, Pf sporozoites reach the liver and establish intracellular infection within host hepatocytes, where they develop during the next 7 days into symptomatic erythrocyte infective merozoites, and later on into transmission driving gametocytes. Developing effective malaria vaccines remains a considerable challenge due to the complex life cycle of Pf, its substantial genetic diversity, and sophisticated immune evasion mechanisms (7, 8). Both cellular and humoral immune responses are associated with protection against malaria, but the precise antigens that, either by themselves or collectively can confer sterile and durable protection are not yet fully known (9, 10). Moreover, the parasite’s large genome, encoding over 5,300 antigens, further complicates identification of key antigens targeted by protective immunity (11, 12), with only a fraction evaluated as vaccine candidates (13). In contrast to antigens expressed during early sporozoite, late merozoite, and gametocyte stages, which were advanced into vaccine designs, liver-stage antigens remain largely uncharacterized, leaving a major blockade in pan-stage vaccine designing approach (14-17).

Immunization with whole sporozoites offers a strong possibility to assess overall protective capacity and discovery of immunogenic antigens. Such approaches utilize distinct attenuation strategies that arrest parasite development in the human host at specific stages. While radiation-attenuated sporozoites (RAS) arrest soon after infecting hepatocytes (18), chemo-attenuation by chloroquine chemoprophylaxis (CPS) allows for full parasite maturation in the liver and subsequent clearance in the blood (19, 20). Recently, we have shown that targeted genetic attenuation arrests the malaria parasites at specific development stages in the liver (21, 22). These genetically attenuated parasites (GAP) offer greater antigenic exposure compared to RAS while arresting precisely before the onset of symptomatic blood stage parasites. We demonstrated that malaria parasites genetically attenuated to arrest late, but not early, during the liver stage development show high protective capacity in malaria naïve humans (22).

In this study, sterile protection was achieved in 8 of 9 participants immunized with a late-arresting GA2 (*PfΔmei2*) parasite, compared to 1 of 8 participants immunized with an early-arresting GA1 (*PfΔb9Δslarp*) parasite (22). Progress in high-density whole-proteome microarrays allows for rapid analysis of humoral responses to infectious diseases, supporting both vaccine antigen identification and serodiagnostic antigen discovery (23-25). In this study, we made use of Pf proteome microarray coated with the selected set of Pf antigens reported to associate with protection in previous studies (26). Using the unique clinical trial samples, here we report the distinct immunogenicity of GA1 and GA2 parasites by analyzing the humoral antibody and cellular responses. Our work uncovered novel antigens targeted by the immune response to late-liver stage arresting parasites and shows that the overall breadth of the humoral immunity correlates with Pf-specific cellular immunity in the clinical trial participants.

## Materials and methods

### Study population design

In the current study (ClinicalTrials.gov registration number NCT04577066), 20 healthy malaria-naïve Dutch participants received bites from 50 mosquitoes, of which 8 participants received bites infected with early arresting GA1 parasites, 9 participants received bites infected with late arresting GA2 parasites, and 3 participants received uninfected bites (22). Participants were immunized three times at 4-week intervals. Among the participants, 1 out of 8 who received GA1 showed protection, while 8 out of 9 receiving GA2 were protected, and all 3 placebos were nonprotected (22). Blood and plasma samples were collected 1 day prior to each immunization (I, II, III) and prior to challenge through controlled human malaria infection (CHMI) with wild type (WT) Pf 3D7 (Fig. 1A). Protection is characterized by the absence of detectable parasitemia as measured in Pf qPCR throughout the 21-day daily follow-up period after challenge with WT Pf 3D7.

**Figure 1:**
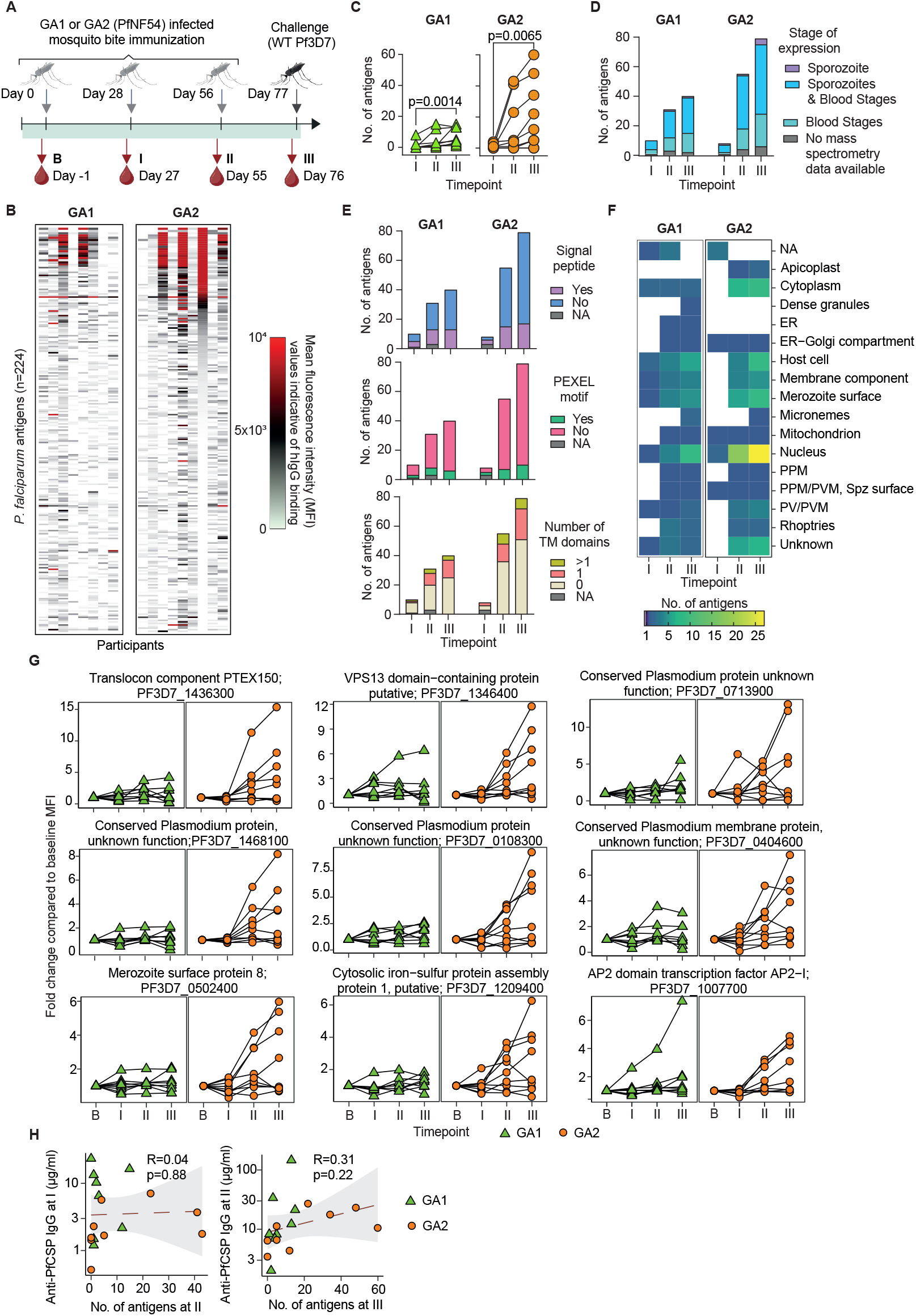
Immunization with the late liver-stage arresting parasite induces a stronger and broader humoral response compared to early-arresting parasite. **A**. A schematic overview of the clinical trial where participants were immunized three times with mosquito bites (MB) infected with either early (GA1) or late (GA2) arresting Pf parasites, followed by a challenge with wildtype (WT) Pf (22). **B**. IgG binding values in plasma collected at (III) from microarray coated with 262 peptides covering 224 unique antigens. **C**. The number of targeted antigens in GA1-MB and GA2-MB participants across three timepoints. Statistical analysis was performed by the Friedman test. **D**. Predicted expression stage of antigens targeted upon GA1-MB and GA2-MB immunization. **E**. Predicted the presence of signal peptide (top), Pf-specific PEXEL motifs (middle), and the number of transmembrane domains (bottom) among the antigens targeted upon GA1-MB and GA2-MB immunization. **F**. Predicted location of targeted antigens after GA1-MB and GA2-MB. **G**. The MFI fold change compared to baseline (B) of Pf antigens with higher seroprevalence in GA2-MB compared to GA1-MB following three immunizations. **H**. Pearson correlations between targeted antigen counts at the indicated immunization timepoint and α-PfCSP antibody titers following the previous immunization.

### Construction and probing of the microarray

A selected Pf3D7 proteome microarray was constructed at University of California, Irvine, CA. Sequence-verified polypeptides were printed as *in vitro* transcription and translation (IVTT) reactions, following previously established methods (27). The T7 promoter plasmids used for encoding each protein in the IVTT reactions were acquired through a high-throughput cloning technique (19). Quality control of the array slides was carried out as described in earlier studies (25), yielding over 97% protein expression efficiency from the spotted IVTT reactions.

Microarray slides were first blocked with the protein array blocking buffer (GVS S.p.A., Zola Predosa, Italy) for 30 minutes. Plasma samples were diluted 1:100 in protein array blocking buffer with an added 10% (vol/vol) *E. coli* BL21(DE3) lysate (Genscript, Piscataway, NJ) and were incubated at room temperature for 30 minutes. The lysate was used to block anti-*E. coli* antibodies in the plasma samples, hence minimizing non-specific background binding. Thereafter, microarray slides were incubated with diluted plasma samples overnight at 4 °C.

Antibody binding was detected using an Alexa Fluor® conjugated goat anti-human IgG Fc fragment-specific secondary antibody (Jackson ImmunoResearch, West Grove, PA), diluted 1:200 in blocking buffer. The microarrays were scanned using a TinyArray imager (28), and spot intensities were analyzed using the ProScanArray Express system (PerkinElmer LAS, Waltham, MA), with spot-specific background correction applied.

Raw data from the scans were expressed as the mean pixel intensity within the printed spots, with surrounding background intensities subtracted for each protein or control. Negative control spots, known as IVTT or “no-DNA” controls, consisted of IVTT reaction mixtures without plasmid templates. The average intensity of these control spots was subtracted from the experimental spots. Correlation analysis of arrays processed on different days and from distinct slide batches confirmed the reproducibility of the slide printing and probing.

### Data normalization, statistical testing and analysis

Data analyses were conducted in the R statistical program (version 4.3.3) and visualized through the ggplot2 R package (29, 30). Additional plots were created in GraphPad Prism (10.2.3), using the built-in statistical tools for analysis. To address variations in baseline antibody levels among immunized individuals and to measure specific antibodies induced by GAP immunization, a computational method implemented with the R package mixtools (31), was used to define binding thresholds for individual participants. The two-component mixture model was developed using a participant’s bound antibody levels at both the baseline (B) and after three immunization (III) timepoints across all 262 peptides. This model differentiates between positive and negative values based on the bound antibody distributions, by means of defining the cutoff point set as the mean of the negative antibody signal distribution plus 3 standard deviations as applied in the previous study (32). Additionally, seropositivity for Pf antigens was considered only when the bound antibody levels at a given timepoint post-immunization were at least two-fold or higher compared to that of baseline.

### Pf-specific cellular immunity measurement

Cellular immunity was measured as described previously (22). In brief, peripheral blood mononuclear cells (PBMCs) isolated from the clinical trial participants were stimulated with Pf-infected and uninfected erythrocytes for 24 h. The cells were stained for specific cell surface markers for central (CCR7^+^CD45RA^-^, T_CM_) and effector (CCR7^+^CD45RA^-^, T_EM_) memory CD4^+^ and Vδ2^+^ γδ T cells. In addition, the cells were stained for intracellular cytokine markers for interferon-γ, tumor necrosis factor-α, interleukin(IL)-2, IL-4, IL-15 and IL-13.

## Results

To investigate the humoral immunity induced against the genetically attenuated parasites that arrest early (GA1) or late (GA2) during the liver stage development, we collected plasma samples at baseline (B), 27 days post first (I) and second (II) immunizations and 20 days post third (III) immunization (Fig. 1A). We used protein microarrays consisting of 262 Pf antigen peptides, representing 224 unique antigens and measured antigen-specific plasma IgG antibodies from the clinical trial participants across timepoints. Heatmaps depicting the bound IgG levels as mean fluorescence intensity (MFI) values indicate the distinct humoral immunity induced by GA1-mosquito bites (MB) and GA2-MB in clinical trial participants (Fig. 1B).

To qualitatively assess the response and account for inter-individual background noise, we pooled the MFI values from the timepoints B and III for each participant separately and defined the threshold for positive signal using a bimodal distribution calculated by mixtools (31), a computational R function (Fig. S1A). By identifying and enumerating Pf antigens with IgG MFI values higher than the threshold, we observed an increase in the total number of antigens targeted over three Pf exposures (Fig. 1C). Notably, humoral immunity to GA2-MB targeted more antigens compared to GA1-MB. As GA1 and GA2 parasites differ in how far they progress into the liver stage development, we analyzed the expression stages of the antigens targeted by humoral immunity. Using the tandem mass spectrometry information available for the antigens from PlasmoDB (11), we found that a large proportion of antigens were expressed during the sporozoite and blood stages (Fig. 1D). However, GA2-MB immunization targeted a higher number of antigens expressed either exclusively in the blood stage or also during the sporozoite stage over three Pf exposures compared to GA1-MB.

To understand whether the antibody response preferentially targeted antigens that are secreted to the parasite periphery, we investigated the presence of signal peptides using PlasmoDB and SignalP6.0 (Fig. 1E) (11, 33). In addition, we performed screening for *Plasmodium* export element (PEXEL) motifs that may signify potential for protein translocation to or beyond the parasitophorous vacuole membrane and thus may indicate localization at the parasite periphery or host cell cytoplasm. Furthermore, we assessed the presence of transmembrane domains in the antigens to predict their localization on the parasite surface, PVM, secretory network vesicles or host membranes. Interestingly, most antigens targeted by the response against GA1-MB and GA2-MB did not possess the signatures for secretion and membrane localization, suggesting an intracellular localization profile. Notably, utilizing PlasmoDB and GO terminology, we found that the majority of antigens were predicted to have nuclear or cytoplasmic localization, whereas a subset of membrane-bound antigens were targeted (Fig. 1F).

To identify response characteristics influenced by the parasite attenuation stage, we estimated the seroprevalence of antigens during GA1-MB and GA2-MB immunizations and compared to the previously reported seroprevalence in naïve individuals upon three WT-MB under CPS (22, 32) (Fig. S1B). The targeted antigens and their seroprevalence in GA2-MB resembled that of CPS more than GA1-MB, suggesting an overlap in antigens expressed and exposed to the immune system in GA2 and WT parasites during the late liver stage. In spite of the liver stage arresting phenotype, we observed several blood stage antigens being targeted by the GA2-MB response, but not by the GA1-MB response. This indicates that these antigens are expressed in both the late-liver stage and the blood stage. We identified key Pf antigens that were preferentially targeted during GA2-MB in comparison to GA1-MB such as translocon component PTEX150 (PF3D7_1436300), the VPS13 domain-containing protein, putative (PF3D7_1346400), and the conserved *Plasmodium* protein (PF3D7_0713900; Fig. 1G). Although the vast majority of the targeted antigens have a predicted nuclear or cytoplasmic localization, some of the key antigens with high seroprevalence in GA2-MB showed membrane localization, indicating the distinct feature of response against late liver stage arresting GA2 parasites. Indeed, we observed a clear booster effect in the IgG antibody levels against these antigens upon three GA2-MB but not GA1-MB, suggesting the formation and reactivation of memory B cells against the antigens. It is also noteworthy that 27 out of the 85 targeted antigens are *Plasmodium* proteins with unknown functions, 23 of which are conserved, highlighting the critical need to better understand their roles in the liver stage development. Additionally, 29 of the 85 targeted antigens are classified as putative, suggesting their function and role during liver-stage development are not accurately annotated. To address the heterogeneity of the GA2-MB response, we asked whether having strong anti-sporozoite antibody levels would limit the response against liver stage antigens. However, we observed no correlation between the pre-MB anti-Pf circumsporozoite protein (PfCSP) antibody levels and the number of antigens targeted post-MB during both second and third immunization, suggesting the observed GA2-MB response heterogeneity is independent of anti-sporozoite immunity (Fig. 1H). Overall, our results show that GA2-MB induced stronger and broader levels of antibodies targeting several late-liver and blood stage antigens compared to GA1-MB. While the majority of the targeted antigens are predicted to be expressed intracellularly, a subset of membrane-bound antigens were also targeted by GA2-MB response.

We previously reported the stronger cellular immunity of GA2-MB compared to GA1-MB by measuring the Pf-specific CD4^+^ and Vδ2^+^ γδ T cells upon stimulating the PBMCs with Pf-infected and uninfected erythrocytes (Fig. 2A) (22). By comparing the cellular immunity at baseline and after three exposures, we observed that both GA1- and GA2-MBs induced monofunctional and polyfunctional T cells that express single or more than one pro-inflammatory cytokines (interferon-γ, tumor necrosis factor-α and interleukin-2), respectively (Fig. 2B). While polyfunctional CD4^+^ T cells are significantly induced by both GA1- and GA2-MBs, only GA2-MB induced polyfunctional Vδ2^+^ γδ T cells.

**Figure 2:**
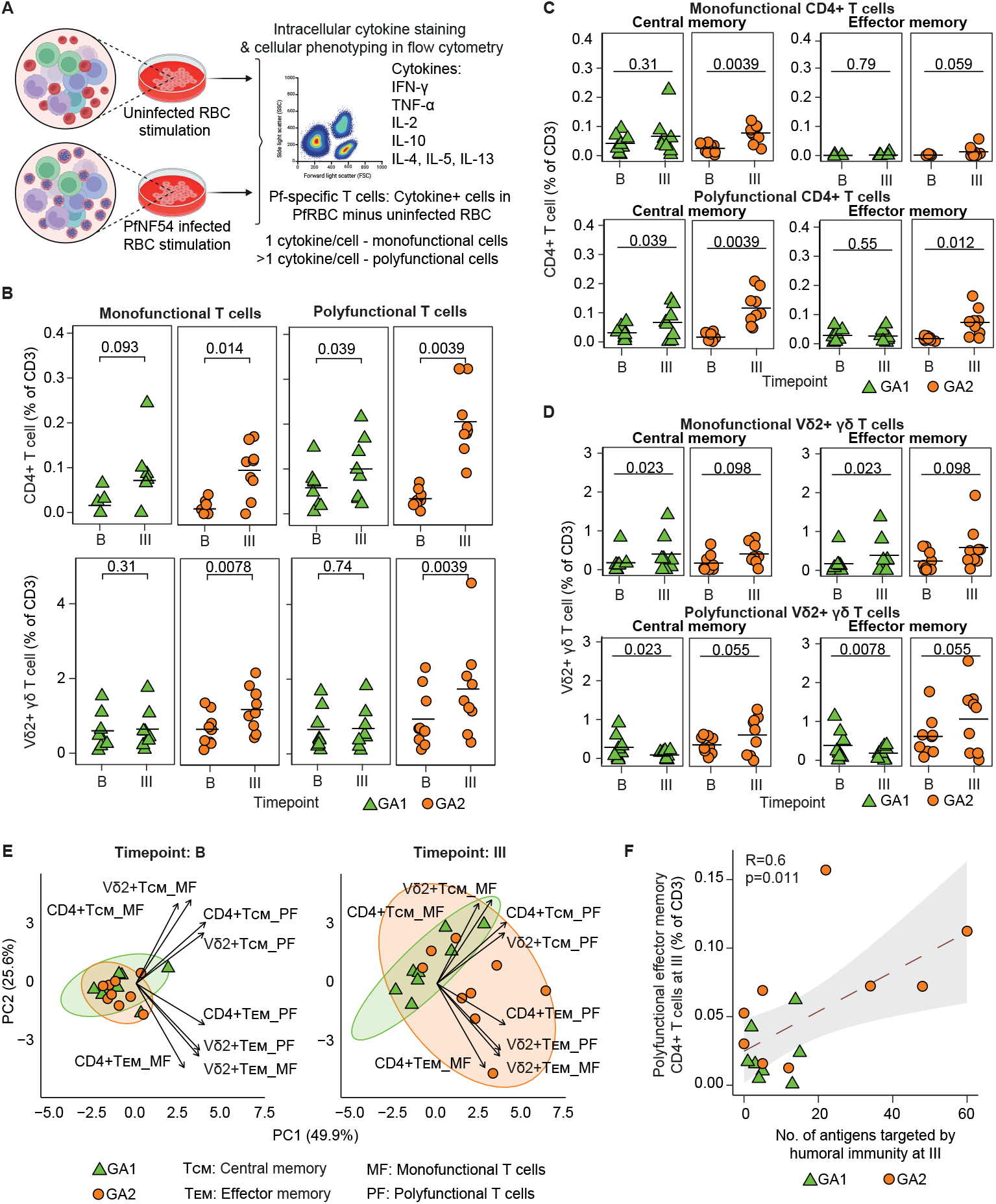
Pf-specific polyfunctional effector memory T cell response correlates with the humoral response. **A**. Experimental design of stimulating PBMCs with Pf-infected or uninfected erythrocytes to detect Pf-specific cellular responses in immunization groups. **B**. Frequency of monofunctional (1 cytokine/cell) and polyfunctional (>1 cytokine/cell) CD4^+^ and Vδ2^+^ γδ T cells after three immunizations (III) compared to the baseline (B). **C-D**. Frequencies of cellular responses defined as central (CCR7^+^CD45RA^-^, T_CM_) and effector (CCR7^-^CD45RA^-^, T_EM_) memory T cells as previously reported among CD4^+^ (C) and Vδ2^+^ γδ (D) T cells (22). **E**. Principal component analysis of mono- and polyfunctional CD4^+^ and Vδ2^+^ γδ T_CM_ and T_EM_ frequencies at both baseline and following three immunizations. **F**. Pearson correlation of polyfunctional CD4^+^ T_EM_ with the number of targeted antigens following three immunizations. B-D Statistical analysis was performed by non-parametric paired t-test.

To identify the distinguishing features of cellular responses, we defined the phenotypic status of Pf-specific T cells as T_CM_ (CCR7^+^CD45RA^-^) and T_EM_ (CCR7^-^CD45RA^-^) as previously reported (Fig. 2C-D) (22). Principal component analysis based on the frequencies of Pf-specific T cells indicated that the effector memory compartment in both CD4^+^ and Vδ2^+^ γδ T cells best distinguished the GA2-MB induced cellular immunity from that of GA1-MB, independent of mono- and polyfunctional status of the responding cells (Fig. 2E). To integrate our findings from the cellular immunity with humoral response measured at the level of individual participants, we compared the frequencies of Pf-specific T cells with the total number of antigens targeted at timepoint III (Fig. 2F, S2A-B). Our analysis revealed the positive correlation between polyfunctional effector memory CD4^+^ T cells and the number of antigens targeted, suggesting the overall coherence between cellular and humoral response in whole sporozoite immunization, particularly when arrested during late-liver stage development.

## Discussion

Once the Pf sporozoite invades a human hepatocyte, the parasite undergoes full maturation inside the infected hepatocyte until the hepatocyte ruptures and releases as mature merozoites into the bloodstream. Hence, protection during the liver-stage development most likely is mediated by cellular but not humoral immunity. However, due to the lack of knowledge of precise antigens expressed during liver stage development in Pf, key antigens that drive the immunogenicity and protective capacity during the parasite liver stage remain unknown (34-36). By studying the humoral response, we here report a list of antigens that are preferentially targeted by the B cells following GA1-MB and GA2-MB immunization. Given the stronger overlap of antigens targeted by GA2-MB with CPS than GA1-MB, the antigens we identified most likely are expressed during the late-but not early-liver stage of the parasite development. These antigens are likely exposed to the immune system either directly during hepatocyte rupture or through acquisition by antigen-presenting cells such as Kupffer cells (37, 38). Furthermore, comparable protective levels achieved by GA2-MB and CPS clinical studies indicate that the response to antigens exposed during the late-liver stage development sufficiently captures the protective response induced in CPS studies (19, 20).

The parasite liver stage development remains a black box as to the antigens that are specifically expressed during this stage, host-parasite interactive elements that are essential for parasite survival and development, and importantly, the antigens that are likely exposed to and targeted by the immune system to establish sterile protection. Studies in murine models are limited by the difference in development duration and antigen repertoire present between murine parasites such as *P. berghei* and *P. yoelii* and human parasite *P. falciparum* (39-41). Nevertheless, these studies indicate the prominent role of monocyte-derived CD11c^+^ cells in acquiring infected hepatocytes and priming liver-stage antigen-specific CD8^+^ T cells (42). However, the parallel role of such innate activation to trigger B cell response is not reported. Given the stronger and broader antibody response against Pf antigens in GA2-MB in comparison to GA1-MB, we believe that the antigen-presenting cells such as previously reported CD11c^+^ cells trigger B cell responses in the hepatic draining lymph node following the acquisition of infected hepatocytes. This would also explain the stronger polyfunctional CD4^+^ T cell response in GA2-MB compared to GA1-MB when stimulated with the PfRBC lysates, a surrogate for late liver-stage antigens. However, whether the antigens targeted by the humoral response elicited polyfunctional CD4^+^ T cells remains to be studied. Future studies should address the potential of the antigens reported here to elicit liver-resident CD4^+^ and CD8^+^ cellular immunity and their capacity to establish protection against malaria.

As superior protective levels of GA2-MB compared to GA1-MB are based on the immunogenicity of the parasite resulting from late-liver stage development, the immunogenic antigens that we report here provide a starting point to study the potential of liver-stage antigens in eliciting cellular immunity and assess their protective efficacy. Current malaria vaccine development approaches are based on inducing high levels of protective immunity against sporozoites (PfCSP) (2, 4), blood stage (Rh5, MSP1) (17, 43) and gametocyte stage (Pfs48/45, Pfs25 and Pfs230) (44). Studies combining Pb sporozoites with liver stage antigens show superior protective capacity in murine models upon challenge with Pb (45, 46). However, whether the orthologue Pf versions of the Pb liver stage antigens induce similar protective capacity remains unknown. Our study identifies a novel collection of antigens that are exclusively targeted by the humoral response of late-liver stage arresting parasite, hence opening new directions for liver-stage targeted subunit malaria vaccine developments.

## Acknowledgements

The authors would like to acknowledge the clinical trial participants for their contribution to this study. The trial received support from the Bontius Foundation. This research has received funding from ZonMw under VIDI grant No. 09150172010035, the European Union under ERC St grant agreement No. 101075876 and Gratama-Stichting/Leids Universiteits Fonds No. 2023-09/W233054-2. RM was supported by the European Union’s Horizon 2020 research and innovation programme under the Marie Skłodowska-Curie grant (101109084).

## Figure legends

**Figure S1:**
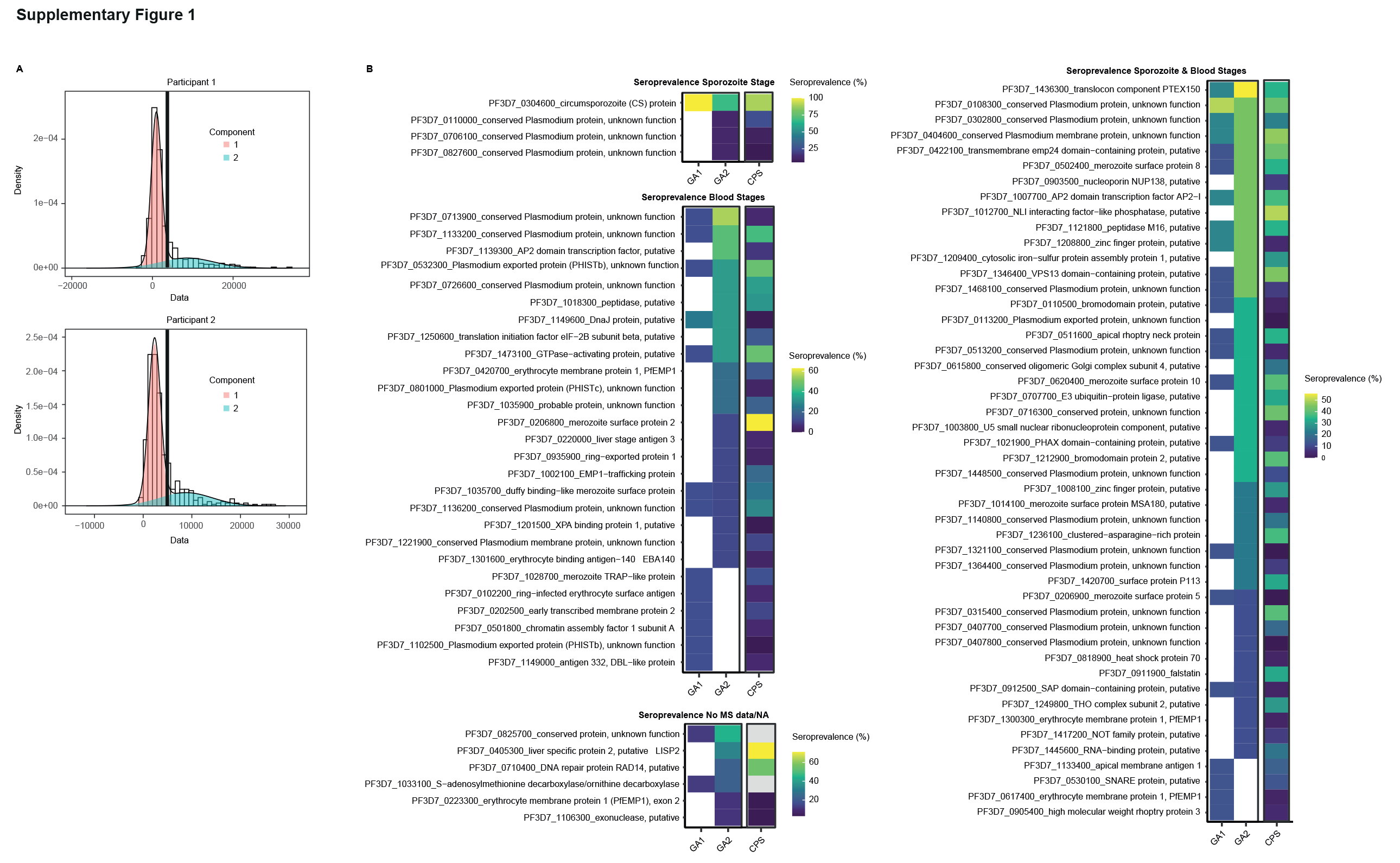
Microarray results representing seroprevalence of Pf-antigens targeted by different immunizations. **A**. Representative bimodal distribution plots of two clinical trial participants indicating negative and positive peaks and the dark black line indicates the threshold for antigen-positivity. **B**. Calculated seroprevalence of Pf antigens in GA1-MB and GA2-MB groups compared with CPS immunized participants published in Oberio et al (32).

**Figure S2:**
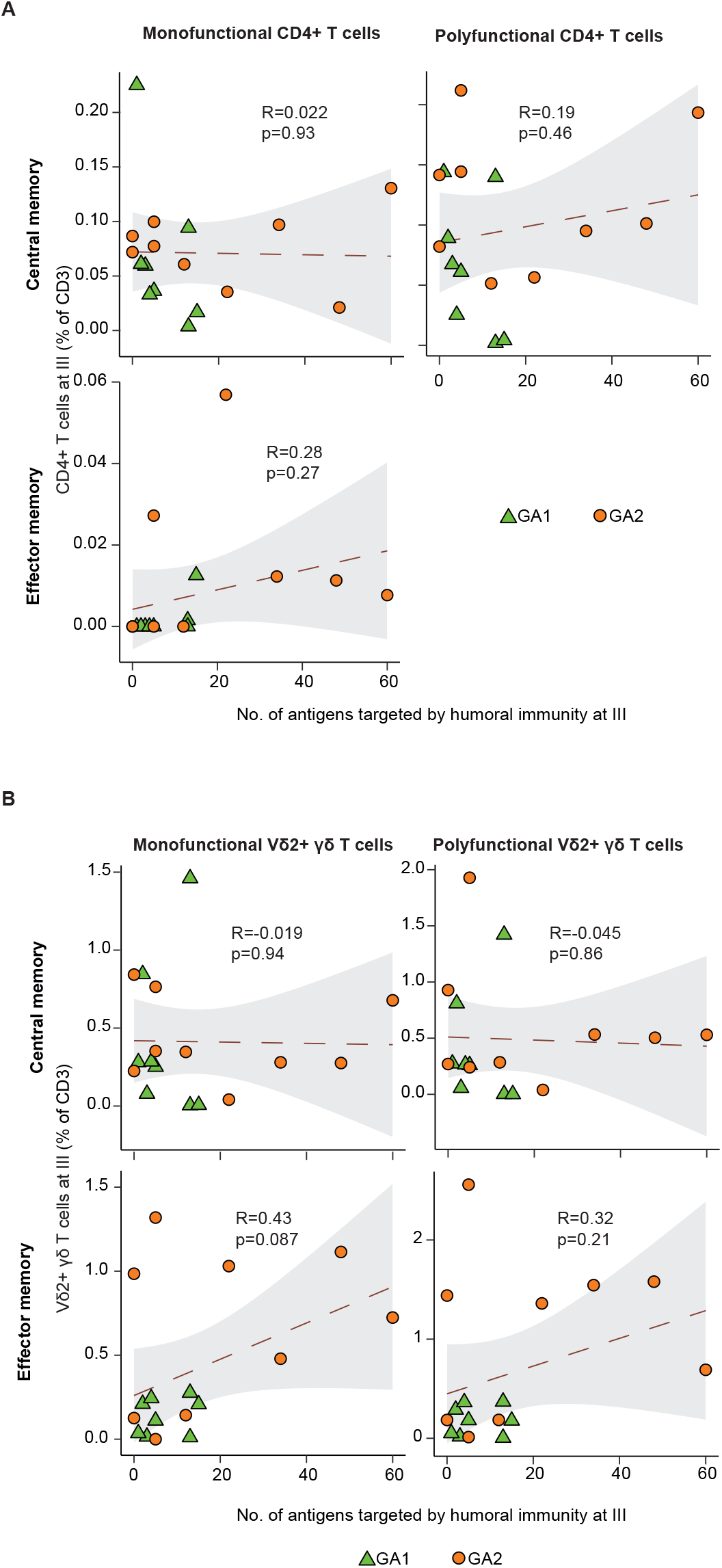
Correlation of Pf-specific cellular response to humoral immunity. **A**-**B**. Pearson correlation of mono- and polyfunctional CD4^+^ (A) and Vδ2^+^ γδ (B) T_CM_ (top) and T_EM_ (bottom) frequencies with the number of targeted antigens following three immunizations.

